# Fuzzy Integral With Particle Swarm Optimization for CNN-Based Motor-Imagery EEG Classification

**DOI:** 10.1101/2021.09.07.459275

**Authors:** Jian-Xue Huang, Ya-Lin Huang, Chia-Ying Hsieh, Chun-Shu Wei

## Abstract

Recently, decoding human electroencephalographic (EEG) data using convolutional neural network (CNN) has driven the state-of-the-art recognition of motor-imagery EEG patterns for brain-computer interfacing (BCI). While a variety of CNN models have been used to classify motor-imagery EEG data, it is unclear if aggregating an ensemble of heterogeneous CNN models could further enhance the classification performance. To integrate the outputs of ensemble classifiers, this work utilizes fuzzy integral with particle swarm optimization (PSO) to estimate optimal confidence levels assigned to classifiers. The proposed framework aggregates CNN classifiers and fuzzy integral with PSO, achieving robust performance in single-trial classification of motor-imagery EEG data across various CNN model training schemes depending on the scenarios of BCI usage. This proof-of-concept study demonstrates the feasibility of applying fuzzy fusion techniques to enhance CNN-based EEG decoding and benefit practical applications of BCI.

## 1 Introduction

Brain-Computer Interfacing (BCI) [1–3] is a pathway that translates brain activities into meaningful signals, providing an alternative method of communication between humans and machines. Such systems enable a subject to send commands only through brain activities. These signals are decoded to distinguish human’s intentions with a high accuracy and can be used in tasks such as motor imagery [4], translating intentions to motor movement. Due to the low signal-to-noise ratio [5, 6] of EEG signals, artifacts which are irrelevant to tasks often exceed the relevant tasks. How to decode EEG by extracting features from informative data with a high quality has been a point of interest [7–10].

A convolutional neural network (CNN) model is a type of neural network comprises multiple convolutional layers and other network components that has been developed for image recognition and computer vision [11]. As one of the most promising model structure in the revolutionary advancement of artificial intelligence, the use of CNN models extends to other domain including EEG data decoding and analysis. Customized CNN models for EEG decoding have outperformed conventional approaches and significantly reduced the tedious handcrafted feature extraction [12–15]. The EEG-specific CNN models were designed to learn the representations of EEG data in both temporal and spatial domain, where the temporal characteristics present the dynamic of brain activity in time and the spatial characteristics locate brain patterns and assess the spatial distributions. EEGNet [12] is a widely used CNN model for EEG data recognition that has a temporal convolutional kernels in its first layer to perform temporal processing on EEG data. On the other hand, when a CNN model incorporates spatial convolutional kernels in the first layer, it extracts fundamental features through spatial combination across channel locations. A representative CNN model with a spatial first layer is SCCNet [14]. The difference between using a temporal or a spatial first layer in a CNN could lead to different views in learning the EEG data. Despite the comparable model size the prediction performance between EEG-Net and SCCNet, different strategies of data analysis could lie within their distinguishable designs of convolutional layers. However, it is clear if fusing the outputs across types of model could positively leverage their predictions and improve the performance in EEG decoding.

In this study, an aggregated framework with model ensemble is proposed to utilize fuzzy fusion techniques [16–23] for integrating outputs of EEGNet and SCCNet. Particle swarm optimization (PSO) algorithm [24] was applied to enhancing the effectiveness of fuzzy integral techniques. The proposed framework was validate on BCICIV-2a, a public dataset of motor imagery EEG recordings with multiple subjects and sessions [25]. The remaining sections are structured as follows. Section II describes the data, the CNN models, and the fuzzy fusion methods. Section III presents the results with discussions. Section IV concludes the findings and contributions of this work as well as a view of prospective work.

## 2 Materials and Methods

The aggregated fuzzy-fusion framework proposed in this study integrates model ensemble of parallel CNN classifiers and fuzzy integrals with PSO. As illustrated in Fig. 1, the input EEG data first undergo the CNN classifiers (EEGNet and SCCNet), and each classifier generates a series of output probabilities corresponding to the classes. Next, the CNN outputs are integrated in the fuzzy fusion block where fuzzy integral with PSO to harness the degree of uncertainty in the class probabilities to yield a fused decision of class label prediction. The proposed framework consists of two major modules, an ensemble of CNN classifiers and a fuzzy fusion block. Two representative CNN models for EEG classification, EEGNet [12] and SCCNet [14] were employed as the sub-classifiers in the ensemble. In the fuzzy fusion block, we validated the combined use of particle swarm optimization on Seguno/Choquet integral [16–18, 26]. The above-mentioned components in the framework are detailed in the following sub-sections.

**Fig. 1.**
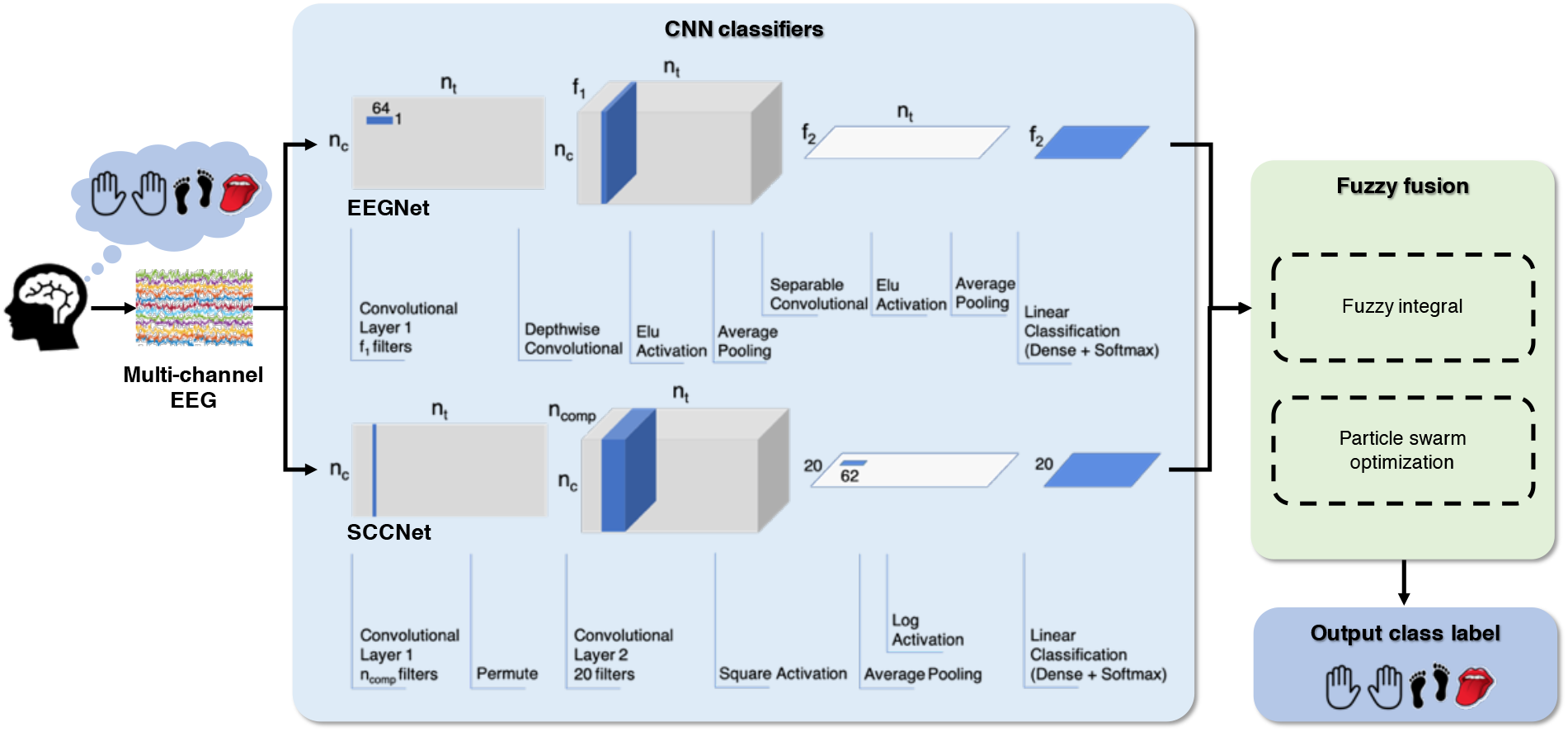
Schematic diagram of the proposed framework comprising: 1) CNN models to convert input EEG data into class probabilities, and 2) Fuzzy fusion using fuzzy integral and particle swarm optimization.

### 2.1 EEGNet

EEGNet is a compact CNN designed specifically for characterizing intrinsic features of EEG [12]. The EEGNet model is able to process multi-channel EEG time series as input, first using a temporal convolutional layer that acts as a series of spectral filters and other temporal operations, followed by a depthwise spatial convolutional layer that serves as spatial filters to enhance the signal and reduce the dimensionality of the data. EEGNet uses separable convolutions to raise the efficiency of the model and maintain a small trainable parameter size. The detail of model architecture is provided as in Table 1. Recent works have shown the capability of EEGNet in learning from relatively small amounts of EEG data across various types of event-related potentials and rhythmic activities [12, 27].

**Table 1.**
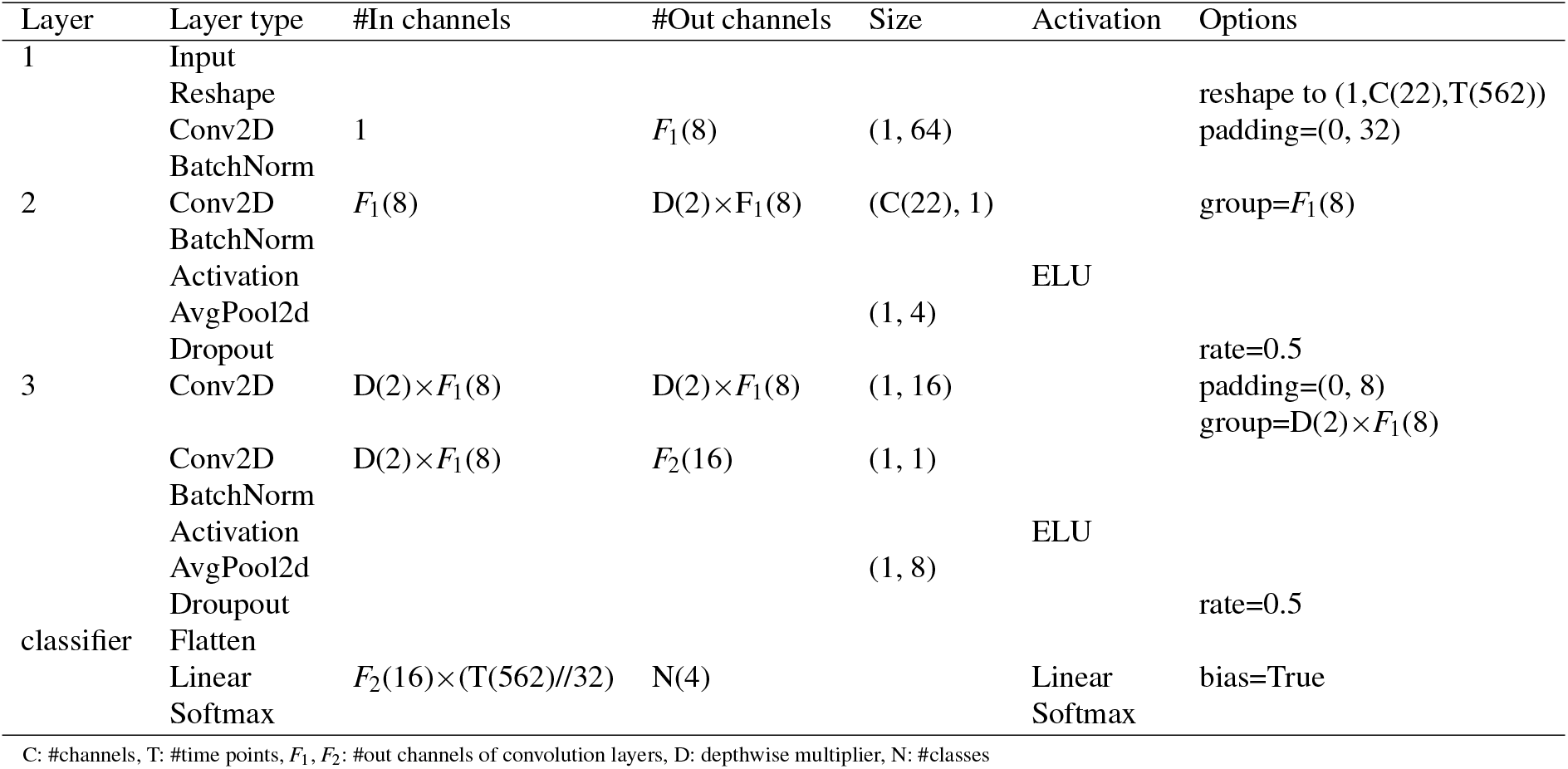
The architecture of EEGNet used in this work.

### 2.2 SCCNet

SCCNet is the abbreviation of ‘Spatial Component-wise Convolutional Network’ [14]. The SCCNet model first process the multi-channel EEG time series using a spatial convolutional layer to filter the data in the spatial domain. This design was inspired from conventional EEG signal processing where spatial filtering techniques (e.g. montage, referencing, component analysis) are often used to reduce noise and to enhance signals in EEG data. In the SCCNet model, a spatial-temporal convolutional layer follows the first spatial convolutional layer to perform temporal filtering and cross relating on the spatial components generated by the first layer. Analogues to EEGNet, SCCNet can learn from relatively small EEG datasets and has a limited model size. The detail of model architecture is summarized as in Table 2.

**Table 2.**
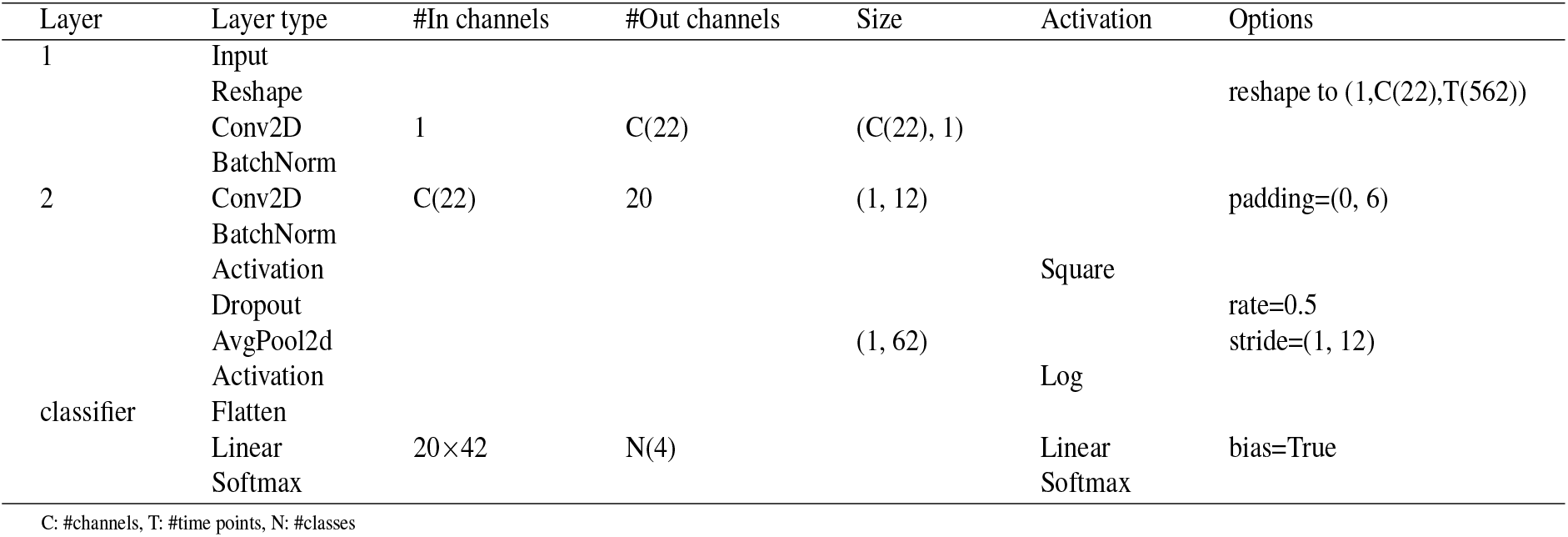
The architecture of SCCNet used in this work.

### 2.3 EEG dataset: BCIC-IV-2a

In this study, we used a well-studied public EEG dataset, the BCI competition IV-2a (BCIC-IV-2a) [25], to evaluate the framework performance. The BCIC-IV-2a dataset consists of motor-imagery (MI) EEG recordings of 9 subjects, and each subject has two sessions recorded on different days. In the MI experiment, a subject was asked to imagine moving different body parts (right hand, left hand, feet, and tongue) for 4 seconds repetitively and thus the number of class is four. Each session contains 72 trials for each class and the EEG data comprise 22 channels of EEG time series. The EEG data pre-processing includes these steps: 1) Downsampling from 256 Hz to 128 Hz, 2) Band-pass filtering at 4-38 Hz, 3) Segmentation by trial at 0.5s-4s of the onset of MI [14].

### 2.4 CNN model training

This section details the information of CNN model training including the division of training/test data and parameter setting in model fitting. Based on the practical scenario of BCI usage, we adopted four training schemes that serves for different conditions of practical BCI usage: 1) Individual training, 2) Subject-independent (SI) training, 3) Subject-dependent (SD) training, and 4) Subject independent training plus fine-tuning (SI+FT) [14]. The four training schemes are depicted below:

- Individual: The individual training scheme refers to dividing training and test data within a single subject. In the BCIC-IV-2a dataset, the first session of a subject served as the training data, and the model was evaluated on the second session of the same subject. This training scheme is applied when a BCI user performs a training session before executing the BCI.
- SI: The SI training scheme is a way to train a model without any information from a target user. This scheme does not require any data from the test subject, but use all of the data from other subjects. For BCI usage, the SI training scheme is able to generate a pre-trained model available for a new user without any training/calibration.
- SD: The SD training scheme incorporates all of the data from other subjects concatenated on the first session of the test subject. This scheme is applicable when existing data are available and the target user has completed a training session before usage.
- SI+FT: The SI+FT training scheme first uses the SI training scheme to train the model, and then applies fine-tuning process on the pre-trained model with first session of the test subject. This scheme utilize exactly the same training data as in the SD training scheme, but it involves two phases of model training rather than a single phases in the SD training scheme. Therefore, the usage of the SI+FT training scheme requires the same condition as for the SD training scheme where a training session is needed for the new user, while a pre-trained model based on all of the data from other subjects should be prepared in advance.

Both CNN models were implemented and evaluated using PyTorch. The model fitting procedure used a batch size of 32, a learning rate of 5 × 10^−4^, and 400 iterations with Adam optimizer. Although the loss function for model fitting was the cross entropy between the predicted class label and the actual class label, the actual output of the CNN models in the context of our proposed framework was a vector of class probabilities generated by the softmax layer.

### 2.5 Fuzzy fusion

Fuzzy integrals utilize the degree of uncertainty in the class probabilities gained from the classifiers [16, 26]. It is regarded as a generalized aggregating operation upon a set of confidence probabilities based on certain continuous weights as fuzzy measures. The fuzzy fusion in the proposed framework is based on fuzzy integral techniques, Segeno integral and Choquet integral, that have previously been successfully applied in various machine learning problems [17, 18].

The Sugeno integral [28–30] is a type of integral with respect to a fuzzy measure. Let *h* : *X* → [0, 1] be an Ω measurable function. The Sugeno integral over a crisp set *A* ⊆ *X* with respect to the fuzzy measure *g* is defined by

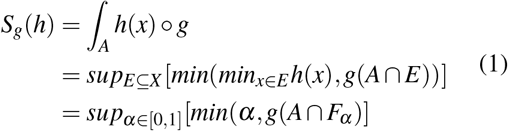

where *F*_*α*_ = {*x*|*h*(*x*) ≥ *α*}

In this study, *A* is equal to {*x*_*i*_, …, *x*_*n*_}, where *x*_*i*_ is the index of model *i. h* refers to the class probabilities generated from the classifiers in the ensemble. *g* is the confidence level of each models.

The Choquet integral [28–30] is a subadditive or superadditive integral created by the French mathematician Gustave Choquet. Suppose *g* is a fuzzy measure, then the Choquet integral of a function *h* : *X* → [0, 1] w.r.t. *g* is defined by

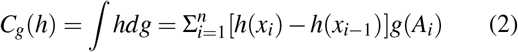

where *A*_*i*_ = {*x*_*i*_, …, *x*_*n*_} and *h*(*x*_0_) = 0.

Here, *h* refers to the class probabilities generated from the classifiers in the ensemble. The other variable, *g*, denotes the confidence level of each models. For Sugeno integral and Choquet integral, the joint confidence [28–30] over a set *A*_*i*_ can be obtained by

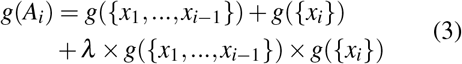

where *λ* can be obtained by solving the equation below:

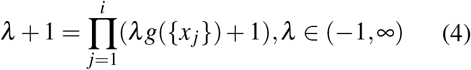

In this work, if the confidence level, *g*, was not determined by the PSO, we assigned *g*(*x*_1_) to 1 and *g*(*x*_2_) to 0.5 as the default setting. We integrated two probability vectors outputted by the two CNN models and got a vector. At the end, the final decision making process for the fuzzy integral fusion yields a vector of modified fuzzy probability corresponding to each class. The final predicted label was determined by the index of the largest value in the vector. In practice, this work employed a open Python implementation of fuzzy integrals available at https://github.com/Fuminides/Fancyaggregations [31–35].

### 2.6 Particle Swarm Optimization (PSO)

Particle swarm optimization (PSO) algorithm is a stochastic optimization method based on swarm. The idea of PSO is to emulate the behaviour of birds or fishes by initializing a set of particles to search for an local or global optima in the desired search-space [28, 29, 36]. Particles are scattered around the search-space, and they move around it to find the position of the optima corresponding to the loss function we defined. It optimizes a problem by iterations to seek a candidate solution with regard to a given measure of quality [28, 37]. To adequately assign a confidence level to the CNN models used in the aggregated framework, the PSO searches for the best confidence level for each CNN model. The PSO algorithm can be defined as following:

1. Randomly initialize the positions for a specific number of particles within the desired search-space.
2. Update the position and velocity of the particles based on the equations:

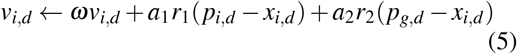

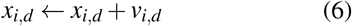

where *ω* is defined as the inertial weight. When *ω* is less than 1, the velocities of all particles will tend to be zero, leading to convergence of the particles into to a local minimum. The coefficients, *a*_1_, *a*_2_, are the acceleration constants, and *r*_1_, *r*_2_ are the random numbers derived from uniform distribution *U* (0, 1). *p*_*g,d*_ is the best known position of the entire search-space in dimension *d*, and *p*_*i,d*_ is the best known position of particle *i* in dimension *d*. Finally, *x*_*i,d*_ stands for the current position of the particle *i* in dimension *d*.

## 3 Results and Discussion

A series of experiment was performed to validate the performance of the proposed fuzzy fusion framework on the motor-imagery EEG data of the BCIC-IV-2a dataset. The fuzzy integral with PSO proposed in this work was evaluated and compared with (i) fuzzy integral without PSO, (ii) simple probability (class decision score) summation, and (iii) individual CNN classification without fuzzy fusion. Meanwhile, our analysis tested the four training schemes for the CNN classifiers based on the context of practical BCI usage. The performance was evaluated by the averaged accuracy across 10 repeats using leave-one-subject-out (LOSO) cross validation. The performances of all fusion techniques with different training schemes are shown in Table 3. The overall performances across 9 subjects of the BCIC-IV-2a were compared using the Wilcoxon signed rank test [38]. Also, the detailed performance of each fusion techniques tested on every subject was provided as in Fig. 2(A)-(D).

**Table 3.**
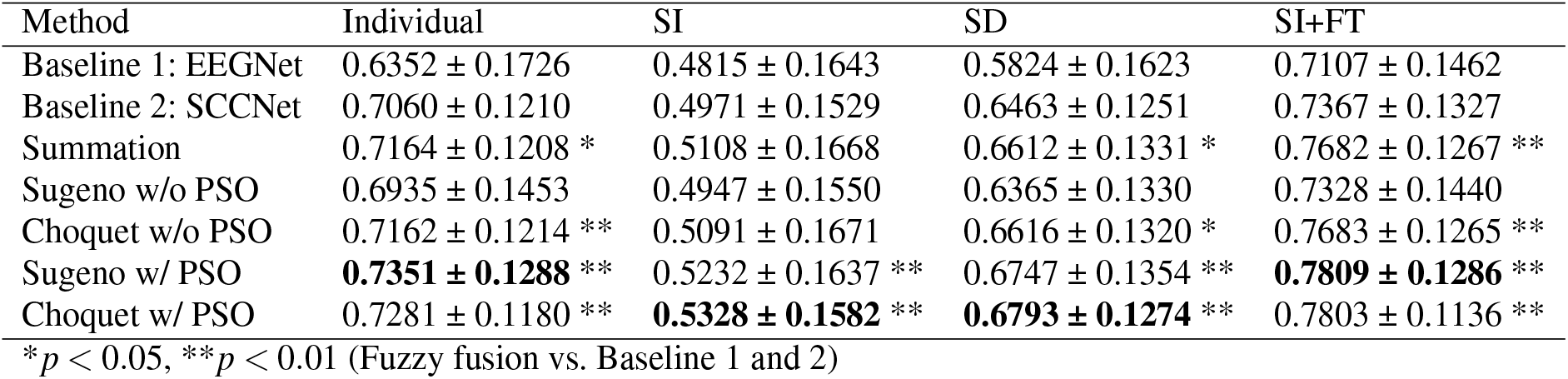
Performance comparison between fusion techniques with various training schemes tested on motor-imagery EEG classification.

| Method | Individual | SI | SD | SI+FT |
| --- | --- | --- | --- | --- |
| Baseline 1: EEGNet | $0.6352 \pm 0.1726$ | $0.4815 \pm 0.1643$ | $0.5824 \pm 0.1623$ | $0.7107 \pm 0.1462$ |
| Baseline 2: SCCNet | $0.7060 \pm 0.1210$ | $0.4971 \pm 0.1529$ | $0.6463 \pm 0.1251$ | $0.7367 \pm 0.1327$ |
| Summation | $0.7164 \pm 0.1208$ * | $0.5108 \pm 0.1668$ | $0.6612 \pm 0.1331$ * | $0.7682 \pm 0.1267$ ** |
| Sugeno w/o PSO | $0.6935 \pm 0.1453$ | $0.4947 \pm 0.1550$ | $0.6365 \pm 0.1330$ | $0.7328 \pm 0.1440$ |
| Choquet w/o PSO | $0.7162 \pm 0.1214$ ** | $0.5091 \pm 0.1671$ | $0.6616 \pm 0.1320$ * | $0.7683 \pm 0.1265$ ** |
| Sugeno w/ PSO | **$0.7351 \pm 0.1288$ \*\*** | $0.5232 \pm 0.1637$ ** | $0.6747 \pm 0.1354$ ** | **$0.7809 \pm 0.1286$ \*\*** |
| Choquet w/ PSO | $0.7281 \pm 0.1180$ ** | **$0.5328 \pm 0.1582$ \*\*** | **$0.6793 \pm 0.1274$ \*\*** | $0.7803 \pm 0.1136$ ** |

\* $p < 0.05$ , \*\* $p < 0.01$ (Fuzzy fusion vs. Baseline 1 and 2)

**Fig. 2.**
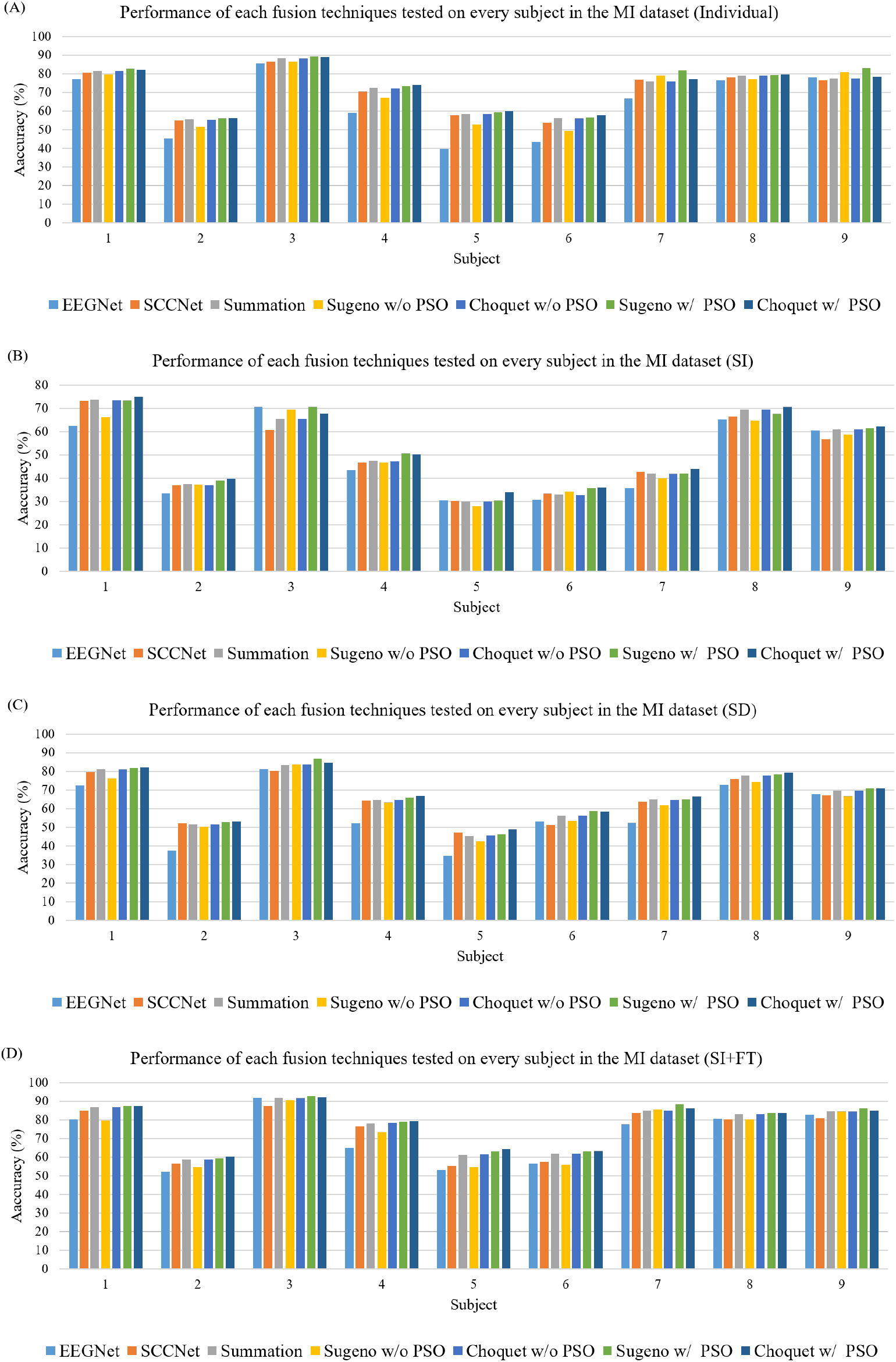
Performance comparison between fusion techniques with different training schemes tested on each subject in the motor-imagery EEG decoding.

As shown in Table 3, fuzzy integral without PSO partially outperform the baseline models. Except for the SI training scheme, simple summation and Choquet integral were able to outperform the baselines in the individual, SD, and SI+FT training schemes without using PSO. When PSO was applied to the fuzzy fusion, the performance was significantly improved with either Sugeno or Choquet integral and achieve the best performances for all four training schemes.

The concepts of the Sugeno integral and the Choquet integral are illustrated in Fig. 3 and Fig. 4 [30, 39, 40], respectively. As shown in Fig. 3, we can divide the output types of the Sugeno integral into four quadrants. When applying PSO to the Sugeno integral, the confidence level of the two CNN models in the ensemble was served by the positions of the particles, which moves around the desired search-space to find the position of the optima. When the value of *g*(*x*_2_) increases, the horizontal red dashed line moves upward. Meanwhile, when the value of *g*(*x*_1_, *x*_2_) increases, the vertical red dashed line moves toward right and expands the area of the rectangle in the lower left corner. Fig. 4 illustrates the behavior of the Choquet integral with two inputs as in our case of two models. The output value of the Choquet integral is the area of the three rectangles. We can simply increase the output value by assigning an increased value of *g*(*x*_2_) or *g*(*x*_1_, *x*_2_).

**Fig. 3.**
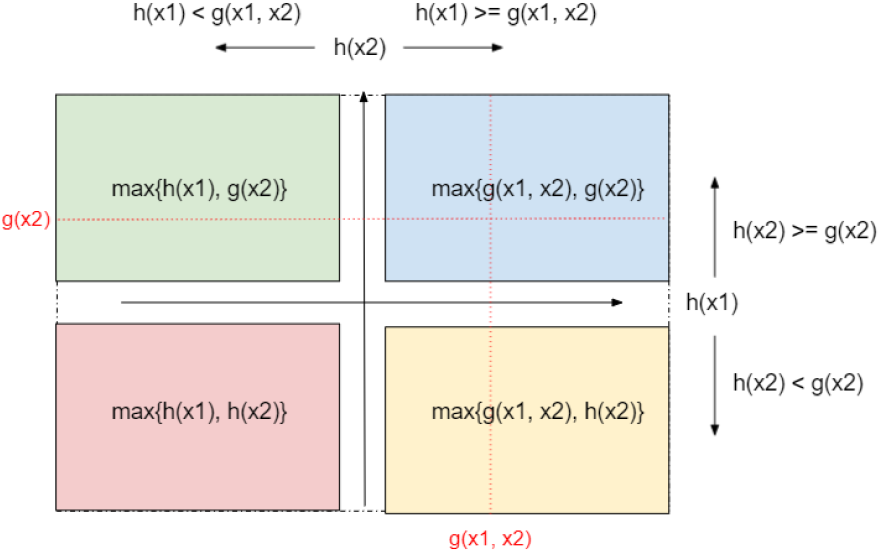
The concept of the Sugeno integral with two inputs.(*h*(*x*_1_) *< h*(*x*_2_))

**Fig. 4.**
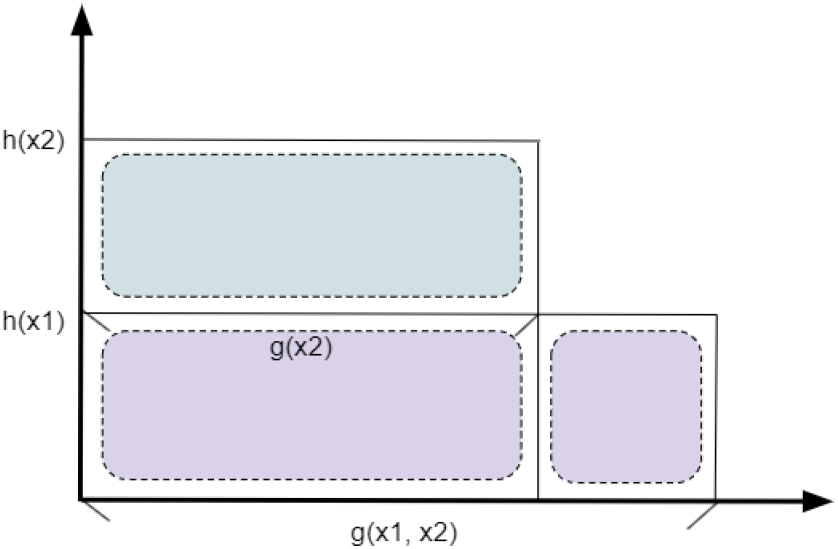
The concept of the Choquet integral with two inputs. [39]

Besides the elevation in the averaged accuracy, the use of PSO led to suppression of standard deviation in the classification performance and thus boosted the robustness of the proposed framework. The PSO algorithm is able to find a proportion which is closed to that best proportion. In general, it allocates a larger area to the output type which achieves a better performance than others. Table 3 exhibits improvements in fuzzy fusion performance after apply PSO. As the CNN models in the ensemble, EEGNet and SCCNet, start the convolutional operation on the EEG data in temporal and spatial domain, respectively, we believe that our fuzzy fusion framework was able to capture the information extracted and enhance the overall performance. Based on the behaviors of the models in the ensemble on a specific session of data, the PSO algorithm automatically tunes the value of *g*(*x*_2_) and *g*(*x*_1_, *x*_2_) (influenced by *g*(*x*_1_) and *g*(*x*_2_)) to distribute the area between four rectangles. There exists a proportion of areas between four rectangles that will maximize the performance of the Sugeno integral. Similarly, the PSO algorithm appears effective in finding an appropriate segmentation between the area of the upper and the lower rectangles as illustrated. The percentage between three rectangles and the total area will decide the performance of the Choquet integral. For both fuzzy integrals, a confidence level in one trial may not be a suitable level in another trial as the optimal confidence level is influenced by the models in the ensemble and the test data. We herein visualize the positions of particles during the iterative process of PSO as shown in Fig. 5. Initially, particles in the search-space are scattered randomly, and gradually shorten the distances to each other while converging to an minimum at the same time.

**Fig. 5.**
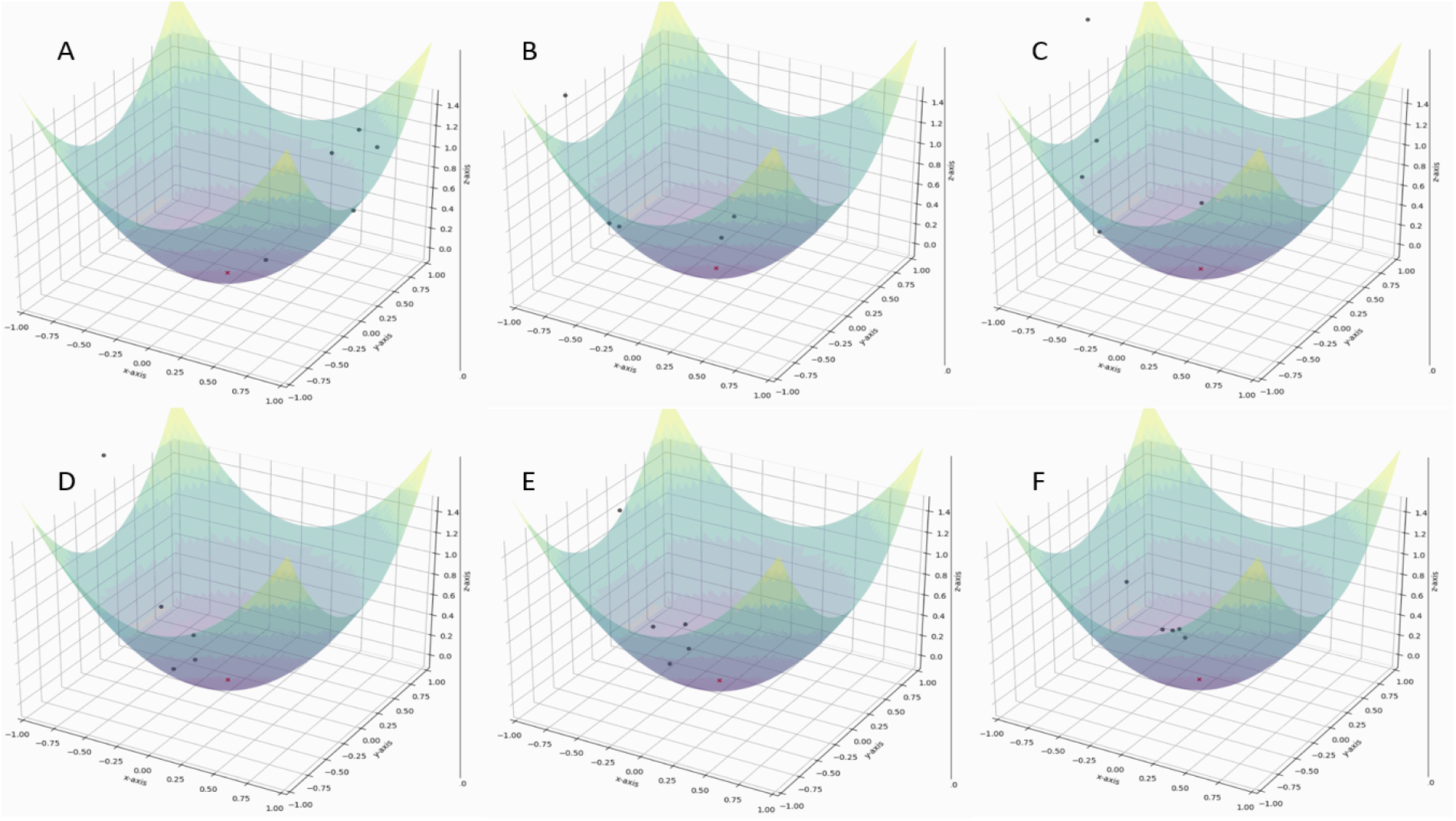
Visualization of a sample run of PSO (the third run testing on Subject 9 using individual training scheme). Dots on the hyper-plane illustrate the dynamic of particle distribution during the training process of PSO with Sugeno integral after (A to F) 1, 10, 20, 50, 80, and 100 iterations.

## 4 Conclusion

In this study, we developed a fuzzy-fusion framework to enhance the performance of EEG decoding beyond individual CNN models (EEGNet and SCCNet). The proposed framework was validated on a four-class motor-imagery EEG dataset across different scenarios of practical applications of BCI. An comprehensive comparison among the fuzzy fusion techniques to validate the performance of our framework under different conditions. The experimental results suggest an partial improvement by using fuzzy integral techniques (Sugeno integral and Choquet integral), as well as a significant enhancement by incorporating the procedure of PSO across model training schemes. This work suggests the feasibility of applying fuzzy fusion to ensemble of heterogeneous CNN classifiers for EEG decoding. As the first proof-of-concept progress in this direction, we believe the use of fuzzy fusion could resolve the issue of selecting a CNN model among different designs for EEG decoding, since the model ensemble is able to accommodate various types of CNN models. We foresee the extension of this work in expanding the model ensemble to a larger number of sub-classifiers and in further evaluations on various types of EEG data in the future.

## 5 Acknowledgment

This work was supported in part by the Ministry of Science and Technology under Contracts 109-2222-E-009-006-MY3, 110-2221-E-A49-130-MY2, and 110-2314-B-037-061; and in part by the Higher Education Sprout Project of the National Chiao Tung University and Ministry of Education of Taiwan. The authors would also like to thank Xin-Yao Huang for his support in implementing CNN models.

## References

[1] J. R. Wolpaw, N. Birbaumer, W. J. Heetderks, D. J. McFarland, P. H. Peckham, G. Schalk, E. Donchin, L. A. Quatrano, C. J. Robinson, T. M. Vaughan et al., “Brain-computer interface technology: a review of the first international meeting,” IEEE transactions on rehabilitation engineering, vol. 8, no. 2, pp. 164–173, 2000.

[2] G. Pfurtscheller, C. Neuper, G. Muller, B. Obermaier, G. Krausz, A. Schlogl, R. Scherer, B. Graimann, C. Keinrath, D. Skliris et al., “Graz-bci: state of the art and clinical applications,” IEEE Transactions on neural systems and rehabilitation engineering, vol. 11, no. 2, pp. 1–4, 2003.

[3] G. E. Fabiani, D. J. McFarland, J. R. Wolpaw, and G. Pfurtscheller, “Conversion of eeg activity into cursor movement by a brain-computer interface (bci),” IEEE transactions on neural systems and rehabilitation engineering, vol. 12, no. 3, pp. 331–338, 2004.

[4] T. Mulder, “Motor imagery and action observation: cognitive tools for rehabilitation,” Journal of neural transmission, vol. 114, no. 10, pp. 1265–1278, 2007.

[5] D. H. Johnson, “Signal-to-noise ratio,” Scholarpedia, vol. 1, no. 12, p. 2088, 2006.

[6] T. Bast, T. Boppel, A. Rupp, I. Harting, K. Hoechstetter, S. Fauser, A. Schulze-Bonhage, M. Scherg et al., “Noninvasive source localization of interictal eeg spikes: effects of signal-to-noise ratio and averaging,” Journal of clinical neurophysiology, vol. 23, no. 6, pp. 487–497, 2006.

[7] V. Bostanov, “Bci competition 2003-data sets ib and iib: feature extraction from eventrelated brain potentials with the continuous wavelet transform and the t-value scalogram,” IEEE Transactions on Biomedical engineering, vol. 51, no. 6, pp. 1057–1061, 2004.

[8] W.-Y. Hsu and Y.-N. Sun, “Eeg-based motor imagery analysis using weighted wavelet transform features,” Journal of neuroscience methods, vol. 176, no. 2, pp. 310–318, 2009.

[9] B. Obermaier, C. Neuper, C. Guger, and G. Pfurtscheller, “Information transfer rate in a five-classes brain-computer interface,” IEEE Transactions on neural systems and rehabilitation engineering, vol. 9, no. 3, pp. 283–288, 2001.

[10] D. P. Burke, S. P. Kelly, P. De Chazal, R. B. Reilly, and C. Finucane, “A parametric feature extraction and classification strategy for braincomputer interfacing,” IEEE Transactions on Neural Systems and Rehabilitation Engineering, vol. 13, no. 1, pp. 12–17, 2005.

[11] K. O’Shea and R. Nash, “An introduction to convolutional neural networks,” arXiv preprint 1511.08458, 2015.

[12] V. J. Lawhern, A. J. Solon, N. R. Waytowich, S. M. Gordon, C. P. Hung, and B. J. Lance, “Eegnet: a compact convolutional neural network for eeg-based brain–computer interfaces,” Journal of neural engineering, vol. 15, no. 5, p. 056013, 2018.

[13] R. T. Schirrmeister, J. T. Springenberg, L. D. J. Fiederer, M. Glasstetter, K. Eggensperger, M. Tangermann, F. Hutter, W. Burgard, and T. Ball, “Deep learning with convolutional neural networks for eeg decoding and visualization,” Human brain mapping, vol. 38, no. 11, pp. 5391–5420, 2017.

[14] C.-S. Wei, T. Koike-Akino, and Y. Wang, “Spatial component-wise convolutional network (sccnet) for motor-imagery eeg classification,” in 2019 9th International IEEE/EMBS Conference on Neural Engineering (NER). IEEE, 2019, pp. 328–331.

[15] A. Krizhevsky, I. Sutskever, and G. E. Hinton, “Imagenet classification with deep convolutional neural networks,” Advances in neural information processing systems, vol. 25, pp. 1097–1105, 2012.

[16] M. F. Anderson, D. T. Anderson, and D. J. Wescott, “Estimation of adult skeletal age-at-death using the sugeno fuzzy integral,” American Journal of Physical Anthropology: The Official Publication of the American Association of Physical Anthropologists, vol. 142, no. 1, pp. 30–41, 2010.

[17] H. Agahi, R. Mesiar, and Y. Ouyang, “On some advanced type inequalities for sugeno integral and t-(s-) evaluators,” Information Sciences, vol. 190, pp. 64–75, 2012.

[18] M. Singh, V. K. Madasu, S. Srivastava, and M. Hanmandlu, “Choquet fuzzy integral based verification of handwritten signatures,” Journal of Intelligent & Fuzzy Systems, vol. 24, no. 1, pp. 145–161, 2013.

[19] G. Salimi-Khorshidi, A. M. Nasrabadi, and M. H. Golpayegani, “Fusion of classic p300 detection methods’ inferences in a framework of fuzzy labels,” Artificial Intelligence in Medicine, vol. 44, no. 3, pp. 247–259, 2008.

[20] X. Luqiang and X. Guangcan, “Study on power spectrum signal fuzzy fusion for motor imagery,” Comput. Eng., vol. 41, pp. 306–309, 2015.

[21] B.-S. Yoo and J.-H. Kim, “Fuzzy integral-based gaze control of a robotic head for human robot interaction,” IEEE transactions on cybernetics, vol. 45, no. 9, pp. 1769–1783, 2014.

[22] L.-W. Ko, Y.-C. Lu, H. Bustince, Y.-C. Chang, Y. Chang, J. Ferandez, Y.-K. Wang, J. A. Sanz, G. P. Dimuro, and C.-T. Lin, “Multimodal fuzzy fusion for enhancing the motor-imagery-based brain computer interface,” IEEE Computational Intelligence Magazine, vol. 14, no. 1, pp. 96–106, 2019.

[23] A. Banerjee, P. K. Singh, and R. Sarkar, “Fuzzy integral based cnn classifier fusion for 3d skeleton action recognition,” IEEE Transactions on Circuits and Systems for Video Technology, 2020.

[24] C. Liu, W.-B. Du, and W.-X. Wang, “Particle swarm optimization with scale-free interactions,” PloS one, vol. 9, no. 5, p. e97822, 2014.

[25] C. Brunner, R. Leeb, G. Müller-Putz, A. Schlögl, and G. Pfurtscheller, “Bci competition 2008– graz data set a,” Institute for Knowledge Discovery (Laboratory of Brain-Computer Interfaces), Graz University of Technology, vol. 16, pp. 1–6, 2008.

[26] M. Grabisch, H. T. Nguyen, and E. A. Walker, Fundamentals of uncertainty calculi with applications to fuzzy inference. Springer Science & Business Media, 2013, vol. 30.

[27] N. Waytowich, V. J. Lawhern, J. O. Garcia, J. Cummings, J. Faller, P. Sajda, and J. M. Vettel, “Compact convolutional neural networks for classification of asynchronous steady-state visual evoked potentials,” Journal of neural engineering, vol. 15, no. 6, p. 066031, 2018.

[28] S.-L. Wu, Y.-T. Liu, T.-Y. Hsieh, Y.-Y. Lin, C.-Y. Chen, C.-H. Chuang, and C.-T. Lin, “Fuzzy integral with particle swarm optimization for a motor-imagery-based brain–computer interface,” IEEE Transactions on Fuzzy Systems, vol. 25, no. 1, pp. 21–28, 2016.

[29] T.-Y. Hsieh, Y.-Y. Lin, Y.-T. Liu, C.-N. Fang, and C.-T. Lin, “Developing a novel multi-fusion brain-computer interface (bci) system with particle swarm optimization for motor imagery task,” in 2015 IEEE International Conference on Fuzzy Systems (FUZZ-IEEE). IEEE, 2015, pp. 1–4.

[30] M. Ayub, “Choquet and sugeno integrals,” 2009.

[31] A. H. Altalhi, J. I. Forcén, M. Pagola, E. Barrenechea, H. Bustince, and Z. Takáč, “Moderate deviation and restricted equivalence functions for measuring similarity between data,” Information Sciences, vol. 501, pp. 19–29, 2019.

[32] H. Bustince, G. Beliakov, G. P. Dimuro, B. Bedregal, and R. Mesiar, “On the definition of penalty functions in data aggregation,” Fuzzy Sets and Systems, vol. 323, pp. 1–18, 2017.

[33] A. Jurio, M. Pagola, R. Mesiar, G. Beliakov, and H. Bustince, “Image magnification using interval information,” IEEE Transactions on Image Processing, vol. 20, no. 11, pp. 3112–3123, 2011.

[34] G. P. Dimuro, G. Lucca, B. Bedregal, R. Mesiar, J. A. Sanz, C.-T. Lin, and H. Bustince, “Generalized cf1f2-integrals: from choquet-like aggregation to ordered directionally monotone functions,” Fuzzy Sets and Systems, vol. 378, pp. 44–67, 2020.

[35] G. Beliakov, H. B. Sola, and T. C. Sánchez, A practical guide to averaging functions. Springer, 2016.

[36] L. J. V. Miranda, “Pyswarms documentation,” 2020.

[37] M. R. Bonyadi and Z. Michalewicz, “Particle swarm optimization for single objective continuous space problems: a review,” Evolutionary computation, vol. 25, no. 1, pp. 1–54, 2017.

[38] R. Woolson, “Wilcoxon signed-rank test,” Wiley encyclopedia of clinical trials, pp. 1–3, 2007.

[39] M.-T. Chu, J. Z. Shyu, G.-H. Tzeng, and R. Khosla, “Using nonadditive fuzzy integral to assess performances of organizational transformation via communities of practice,” IEEE Transactions on Engineering Management, vol. 54, no. 2, pp. 327–339, 2007.

[40] A. J. Pinar, T. C. Havens, M. A. Islam, and D. T. Anderson, “Visualization and learning of the choquet integral with limited training data,” in 2017 IEEE International Conference on Fuzzy Systems (FUZZ-IEEE). IEEE, 2017, pp. 1–6.

